# Classification of Movement-Related Oscillations in sEEG Recordings with Machine Learning

**DOI:** 10.1101/2022.03.28.486094

**Authors:** Alexander P. Rockhill, Alessandra Mantovani, Brittany Stedelin, Ahmed M. Raslan, Nicole C. Swann

## Abstract

Previous electrophysiological research has characterized canonical oscillatory patterns associated with movement mostly from recordings of primary sensorimotor cortex. Less work has attempted to decode movement based on electrophysiological recordings from a broader array of brain areas such as those sampled by stereoelectroencephalography (sEEG). Here we decoded movement using a linear support vector machine (SVM). We were able to accurately classify sEEG spectrograms during a keypress movement in a task versus those during the inter-trial interval. Furthermore, the important time-frequency patterns for this classification recapitulated findings from previous studies that used non-invasive electroencephalography (EEG) and electrocorticography (ECoG) and identified brain regions that were not associated with movement in previous studies. Specifically, we found these previously described patterns: beta (13 - 30 Hz) desynchronization, beta synchronization (rebound), pre-movement alpha (8 - 15 Hz) modulation, a post-movement broadband gamma (60 - 90 Hz) increase and an event-related potential. These oscillatory patterns were newly observed in a wide range of brain areas accessible with sEEG that are not accessible with other electrophysiology recording methods. For example, the presence of beta desynchronization in the frontal lobe was more widespread than previously described, extending outside primary and secondary motor cortices. We provide evidence for a system of putative motor networks that exhibit unique oscillatory patterns by describing the anatomical extent of the movement-related oscillations that were observed most frequently across all sEEG contacts.

**Significance Statement:** Several major motor networks have been previously delineated in humans, however, much less is known about the population-level oscillations that coordinate this neural circuitry, especially in cortex. Therapies that modulate brain circuits to treat movement disorders, such as deep brain stimulation (DBS), or use brain signals to control movement, such as brain-computer interfaces (BCIs), rely on our basic scientific understanding of this movement neural circuitry. In order to bridge this gap, we used stereoelectroencephalography (sEEG) collected in human patients being monitored for epilepsy to assess oscillatory patterns during movement.

## Introduction

Several spectral power changes in electrophysiological recordings related to the initiation of movement have been extensively replicated. Beta power (13 - 30 Hz) decreases (desynchronizes) several hundred milliseconds before movement and subsequently rebounds immediately following the movement. This has been shown most prominently in the subthalamic nucleus and globus pallidus local field potentials (Brown & Williams, 2005) and in sensorimotor electrocorticography (ECoG) recordings (Crone, 1998a; Miller et al., 2007; Stolk et al., 2019). In sensorimotor areas, broadband gamma (60 - 90 Hz) also increases around the time of movement and immediately after (Ball et al., 2008; Crone, 1998b), alpha power has been observed to both increase (Stolk et al., 2019) and decrease/desynchronize (Crone, 1998a) during movement and a robust movement-related negative evoked potential emerges in the time domain 1 - 2 seconds before movement (Toro et al., 1994). These studies have focused on specific brain areas implicated in movement from early electrophysiological and neurosurgical work (Penfield & Boldrey, 1937) and have studied the most prominent oscillations observed in those regions. However, recent work in a rodent model has suggested that movement related activity is highly distributed throughout the brain, perhaps even more so than higher cognitive processes (Steinmetz et al., 2019). Invasive stereo-electroencephalography (sEEG) recordings are well-suited for determining the spatial extent of these movement-related oscillations in humans because they sample more brain areas than electrocorticography (ECoG). In particular, sEEG extends the sampled region to include both deeper cortical structures as well as subcortical regions compared to ECoG. Stereo-EEG also can also record from white matter, and from gray matter without interference from pia and arachnoid mater, which can reduce the signal amplitude (Avanzini et al., 2016). This makes sEEG recordings a promising avenue of inquiry for studying whole-brain oscillatory patterns.

Determining which oscillatory patterns are related to movement is especially suitable for machine learning. Testing each frequency of oscillation at each time point of the epoch surrounding the movement to see if it differs from baseline would cause issues of multiple comparisons. Determining significant time-frequency points using cluster permutations is one way to address this (Maris & Oostenveld, 2007). The cluster permutation method is ideal for comparison with a machine learning approach because it has been validated by simulations and is widely used in electrophysiology research (Pernet et al., 2015). However, cluster permutation cannot account for complex, non-linear effects that are accessible to machine learning approaches. By training a machine learning classifier on seperate data than is used to quantify the accuracies of the classification, more complex patterns can be determined in a way that is repeatable; if the classifier performs well on unseen data, it is likely to perform well on all data recorded with that particular experimental configuration. Thus, a machine learning approach has the potential to quantify complex, higher-order oscillatory patterns that relate to movement.

The ideal machine learning method, in this case, is capable of classification given a limited amount of training data, and is one for which the strength and direction of movement-related time-frequencies can be determined. Linear support vector machines (SVM) are a method of machine learning where a linear hyperplane is iteratively fit to the data to optimally separate the data in different groups (Buitinck et al., 2013). A coefficient matrix that determines the relative importance and direction of each time-frequency pixel in the spectrogram can then be determined from the hyperplane. We chose to use an SVM to classify spectrograms of sEEG recordings during movement because of this interpretability in order to understand which oscillations are related to movement. We hypothesized that event-related potentials and oscillatory patterns would be able to classify periods of movement from inter-trial periods of rest in a broad array of brain areas recorded from with sEEG.

## Methods

### Participants

Eight patients with medically intractable focal epilepsy were surgically implanted with sEEG electrodes for clinical localization of ictal (seizure) onset and epileptogenic zones. All patients underwent robotic assisted 0.8 mm diameter stereoencephalography electrodes (PMT, Chanhassen, MN, USA) with center-to-center pitch of 3.3 to 5 mm for electrode contacts. The total number of electrode contacts analyzed was 979 with 122 +/- 2 contacts per subject. The contacts were distributed in the patients’ brains as shown in *Figure 2*. Patient demographic and clinical information is shown in *Table 1*.

**Table 1.** Patient demographic information and task performance information. Abbreviations: deep brain stimulation (DBS), responsive neurostimulation (RNS), focal cortical dysplasia (FCD).

| ID | Age | Sex | Hand | Response Time (s) | Task Accuracy (Missed trials) | Events Used | Diagnosis of Origin | Surgery Based on sEEG | Notes |
| --- | --- | --- | --- | --- | --- | --- | --- | --- | --- |
| sub-1 | 44 | M | R | 0.447 +/- 0.118 | 98.6% (0) | 202 | Left entorhinal cortex | None | Photodiode displaced for one block |
| sub-2 | 47 | F | L | 0.858 +/- 0.245 | 100.0% (1) | 280 | Bilateral middle and superior temporal gyri | Bilateral anterior thalamic nucleus DBS |  |
| sub-5 | 33 | F | R | 1.373 +/- 0.624 | 99.3% (12) | 291 | Bilateral temporal pole, amygdala, left orbitofrontal | Ablation of right frontal FCD and right temporal pole and bilateral anterior thalamic nucleus DBS |  |
| sub-6 | 31 | F | R | 0.401 +/- 0.058 | 99.0% (0) | 298 | No seizures | Right anterior temporal lobectomy |  |
| sub-9 | 39 | M | R | 0.558 +/- 0.266 | 97.9% (8) | 286 | No seizures | None |  |
| sub-10 | 31 | F | R | 0.615 +/- 0.276 | 94.2% (7) | 289 | Left hippocampus, amygdala, parahippocampal gyrus | Left amygdalo-hippocampectomy |  |
| sub-11 | 30 | M | L | 0.595 +/- 0.485 | 100.0% (0) | 134 | Left amygdala and hippocampus | RNS in bilateral hippocampus | Photodiode displaced for two blocks |
| sub-12 | 26 | M | R | 0.434 +/- 0.118 | 98.3% (0) | 279 | Temporal-frontal neocortical and mesial regions | None | Previous surgical resection of right temporal lobe |

#### Task

Patients performed a forced two-choice reaction time task with the left and right arrow keys on a laptop as shown in *Figure 1*. On each trial, patients were presented with a fixation cue for 300-700 ms and then a left or right arrow was presented. Participants were asked to respond before the arrow disappeared with the corresponding key on the laptop keyboard, with a total of 300 trials. The duration of the cue varied between 1.5 times and 4 times each participant’s average reaction time during 10 practice trials. The task was administered using a custom MATLAB script implementing PsychToolBox (Brainard, 1997; Kleiner et al., 2007; Pelli, 1997). The laptop was placed on the patients’ laps while reclined in a hospital bed. A photodiode was attached to the bottom right corner of the screen and was output to the amplifier to synchronize the task to the electrophysiology. Due to noise in the hospital environment a few photodiode events were corrupted for most patients (see *Table 1*), and, due to the patient displacing the photodiode by shifting in bed, two blocks of 75 trials for one patient and one block of 75 trials for another patient were unusable. Accurate timing could not be derived from trials without photodiode synchronization between the electrophysiology recording and the task so these trials were excluded. Trials with incorrect responses were also excluded. Participants *sub-2* and *sub-11* were left handed and used that hand to perform the task, the rest were right handed and used their right hand for the task.

**Figure 1.**
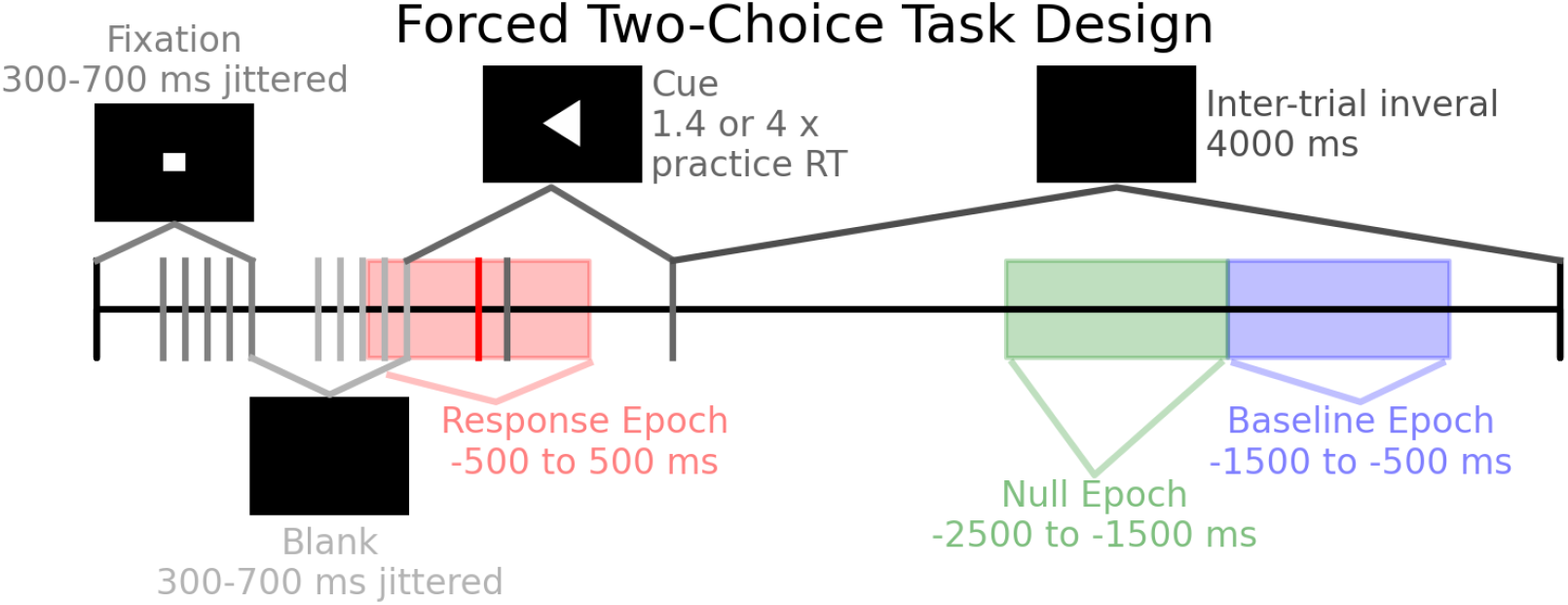
The task schema is shown per trial. The diagram is to scale in time. A typical response time (RT) is shown in red. Trials were included if the response was correct and the response time was within the cue period.

**Figure 2.**
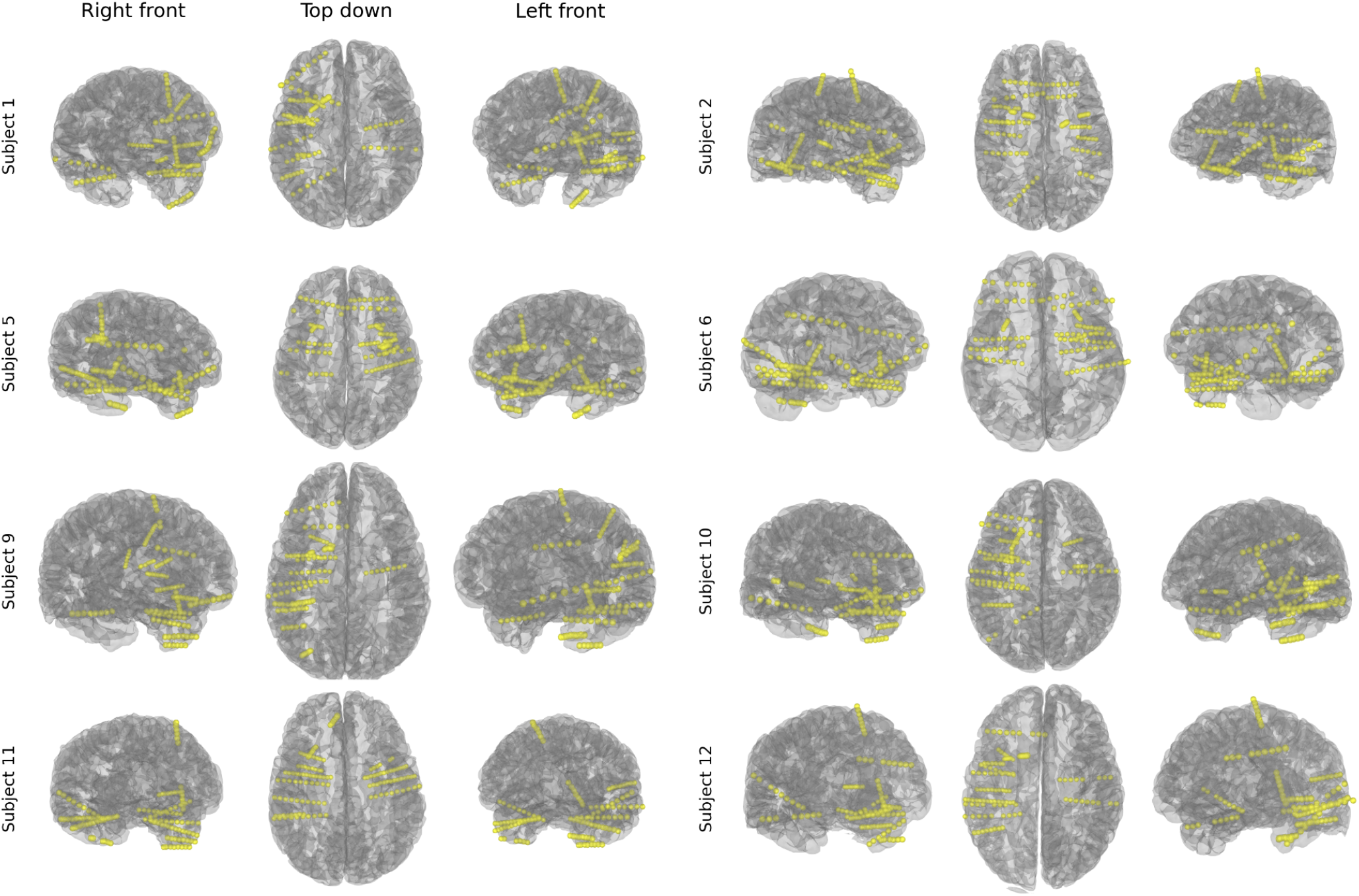
Surgical lead placements for the eight patients shown in a 3D rendering from three perspectives. Coverage was generally biased towards unilaterality and temporal lobe, but a wide range of brain areas were covered.

#### Preprocessing

Electrode positions were determined with MNE-Python (Gramfort, 2013) using the patients’ postoperative computed tomography (CT) image registered to their preoperative magnetic resonance (MR) image (A. Rockhill et al., 2022). Anatomical labels were assigned to contacts based on the Deskian-Killiany atlas label of the Freesurfer reconstruction based on that patient’s individual anatomy (Fischl, 2012). The contact locations were then warped to a template brain (*cvs_avg35_inMNI152*) for comparison of their relative spatial locations. The task-related fixation, cue and response events were synchronized using the photodiode (A. P. Rockhill et al., 2020). The time-frequency spectrogram for each event was computed via the Morlet wavelets method with frequencies from 1 to 250 Hz using MNE-Python (Gramfort, 2013; Harris et al., 2020). The voltage time-series signal, bandpass filtered between 0.1 and 40 Hz, was appended to the bottom of the spectrogram in order to include the event-related potential in the classification.

#### Classification

Movement spectrograms consisted of the period 0.5 seconds before the response key was pressed to 0.5 seconds after. These were classified as different from an equal length spectrogram during the inter-trial interval. First, the training spectrograms were dimensionally reduced via principal component analysis (PCA). The first 50 principle components were used so that most of the variance was captured while still decreasing the dimensionality. The PCA components were used as input to a linear SVM classifier implemented in scikit-learn (Buitinck et al., 2013). An 80 / 20% training-test split was used. In addition, another equal length period during a separate part of the inter-trial interval was used instead of the response period as a null classification as shown in *Figure 1*. The coefficient matrices from the SVM, which showed the correlations of time-frequency points with the movement classification, were validated using a one sample cluster permutation test implemented in MNE-Python (Gramfort, 2013). The threshold was set at 99% of a T distribution (*alpha*=0.01) with degrees of freedom one less than the number of subjects. For each sEEG channel, clusters with T-statistics more extreme than 99% (*alpha*=0.01) of permuted clusters were considered significant.

The SVM method was compared to common spatial pattern (CSP) decoding, a well-used approach in electrophysiology signal classification, for further validation (Gramfort, 2013; Koles et al., 1990). The key difference between the two methods was that the SVM classification was per-contact and so did not use any information about which patient the contact was recording from, whereas the CSP classification was per-patient and used the montage of all contacts implanted in a patient as the basis for classification. For the CSP classification, the voltage signal was bandpass filtered for each frequency and then the spatial pattern was classified using a linear discriminant analysis with 5-fold cross-validation. Thus, the linear SVM was agnostic of the spatial relationship between the sEEG contacts whereas the spatial relationship was the basis of CSP classification; this difference in approach made CSP an ideal validation method.

## Results

The linear SVM using principal components was an effective method of classifying movement spectrograms from spectrograms during the inter-trial interval. The mean of the distribution of scores was unlikely to occur by chance (paired t-test, *p* < 0.001, df = 975) compared to the null distribution generated from the classification with a separate inter-trial interval period instead of the period during movement. At an *alpha*=0.01 relative to the null distribution, 403/975 contacts had classification probabilities that were significant (*Figure 3*). The SVM classification showed that sensorimotor areas had the highest classification accuracy, followed by prefrontal areas (*Figure 4* and in greater detail by specific area in *Figure 5*). Despite having a greater amount of coverage, temporal areas generally had lower accuracy of classifications, but, interestingly, some temporal areas still had contacts with spectrograms that were able to be classified at accuracies that were unlikely to occur by chance.

**Figure 3.**
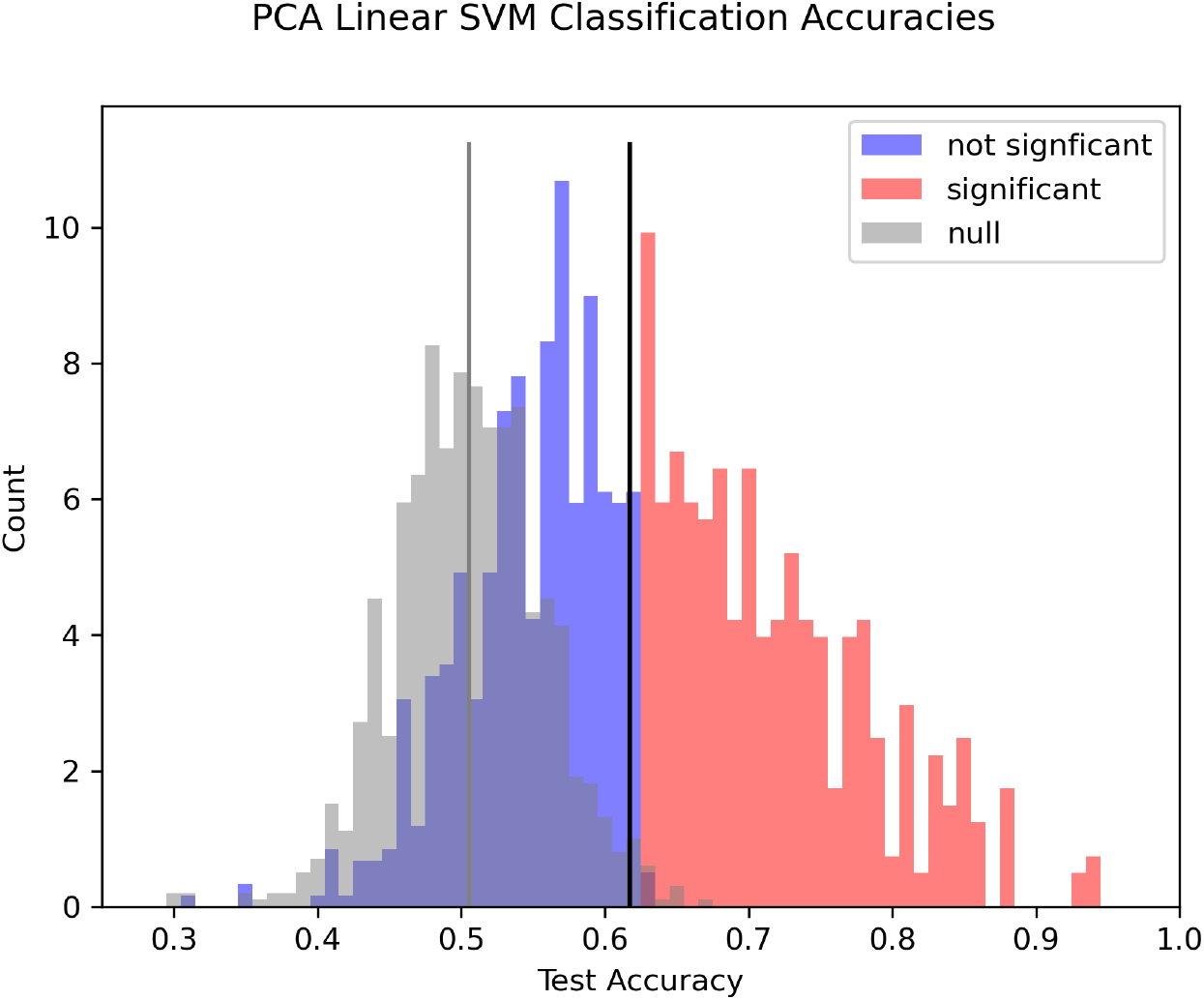
A histogram of the classification accuracy for all electrode contacts across all patients for the SVM classification. Contacts that are significant at an alpha=0.01 level relative to the null distribution for pooled across all contacts are shown in red, those that are not are shown in blue. The mean of the total distribution of scores is indicated with a black line, compared to the mean of the null distribution which is shown in gray (paired t-test, p < 0.001, df = 975). Note that each patient had a different number of trials that were used as shown in Table 1 (see Methods for explanation of missing trials) but the classification accuracies are pooled since accuracy is a normalized metric.

**Figure 4.**
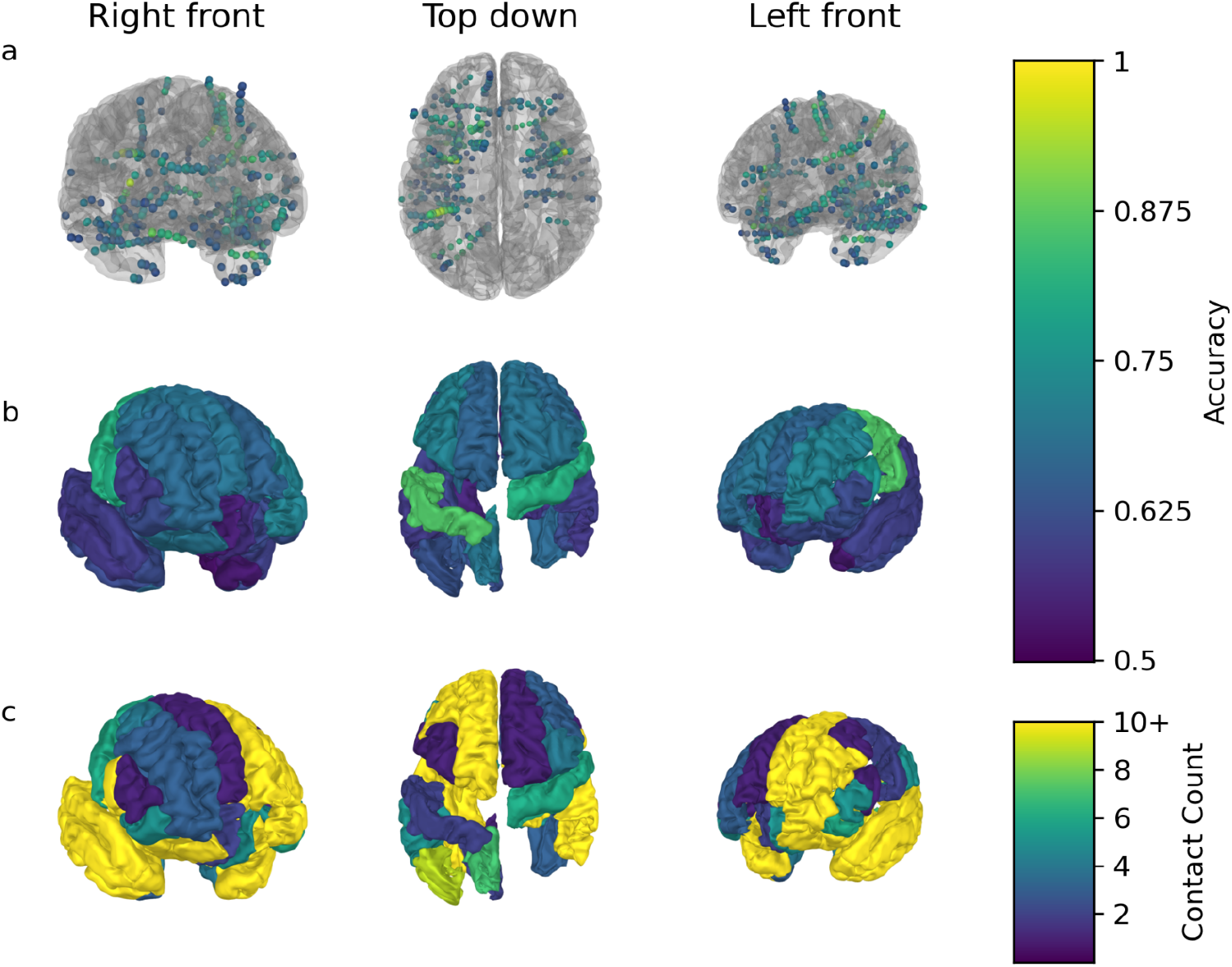
Electrode contacts which had spectrograms that were classified with accuracies significant at an alpha=0.01 level by the SVM are colored by accuracy per contact (a) and averaged per region (b). The number of contacts implanted in each brain region across all patients for this study is shown in c. The highest accuracy was in sensorimotor regions followed by frontal areas with notably lower accuracy in temporal contacts. Regions without data are not shown.

**Figure 5.**
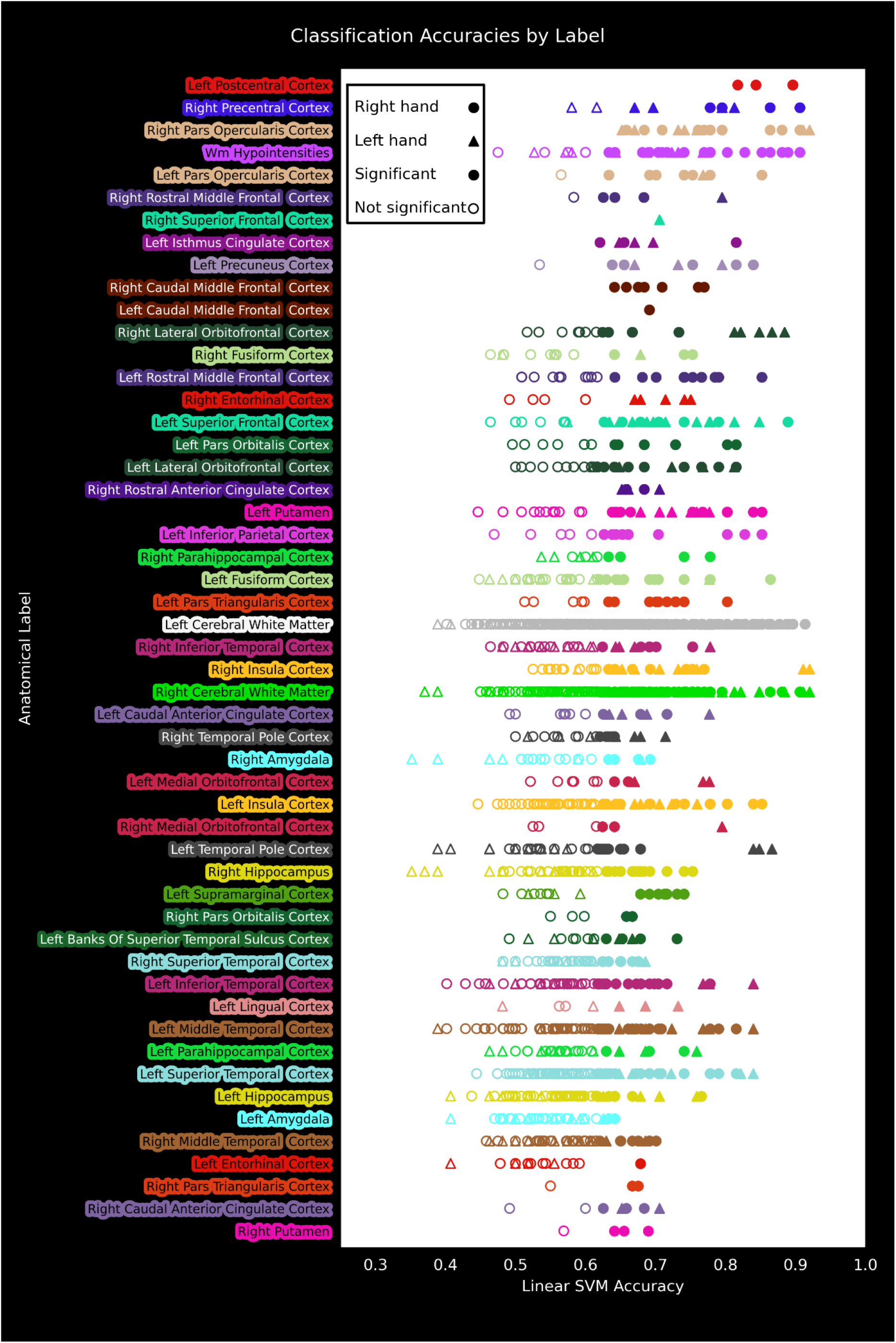
The classification accuracies of the SVM are shown for anatomical labels using the Desikan-Killiany atlas parcellation of the individual patient anatomy. Labels within 3 mm of the center of the electrode contact were assigned to that contact. Note that 22 contacts were unlabelled and are not shown. Some contacts were unlabelled because Freesurfer’s automated labeling process failed near lesions from previous epilepsy surgeries, contacts were above the pial surface (this happened when the deepest contact reached its target when the most superficial contact was still above the pial surface), or limited resolution of the MR and imperfect labeling process could not assign a label.

The CSP analysis showed that beta power was an important classification band in 5/8 patients (*Figure 6*). Additionally, four patients were classified by alpha power, with one patient’s classification depending exclusively on alpha power (Subject 10), suggesting this is also an important oscillatory pattern. Two patients (Subjects 11 and 12) were not able to be classified effectively using this method at all. Notably, the time-frequency features with high classification accuracy in the CSP analysis, were nearly identical to the features from the SVM coefficient matrices and cluster permutation analysis (*Figures 7 and 8*). The CSP classification accuracies also corresponded well to those of the SVM; patients with worse CSP classification accuracy had fewer contacts that had significant classifications. For patients without high-accuracy CSP classifications (Subjects 11 and 12), fewer contacts had significant classification accuracies using the SVM. Overall, the CSP and SVM classifications were concurrent in that most patients had coverage of areas that were modulated by movement. This highlighted how widespread movement-related oscillations are, while, at the same time, suggesting such patterns were confined to specific neural circuitry that was not well-sampled in every patient.

**Figure 6.**
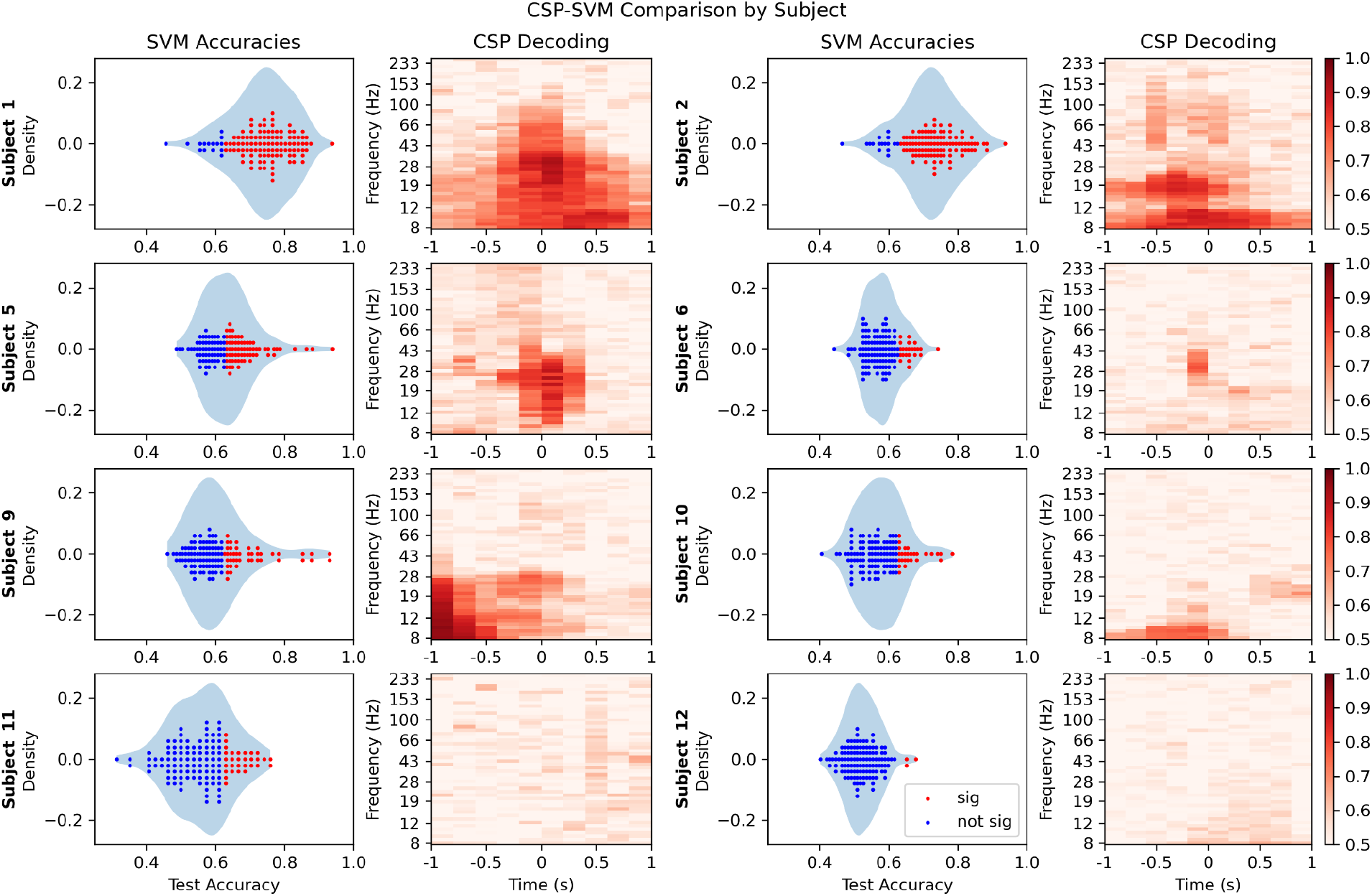
SVM and CSP classifier accuracies are shown on the left and right respectively of each pair of plots per patient. On the left, channels with significant (alpha=0.01) classifications relative to the null distribution for that patient are plotted in red, over a density plot of the distribution. On the right is the classification accuracy using the spatial pattern at each time-frequency bin. Note that patients with more significant classification channels (shown in red on the left) had greater CSP accuracies (darker red on the right).

**Figure 7.**
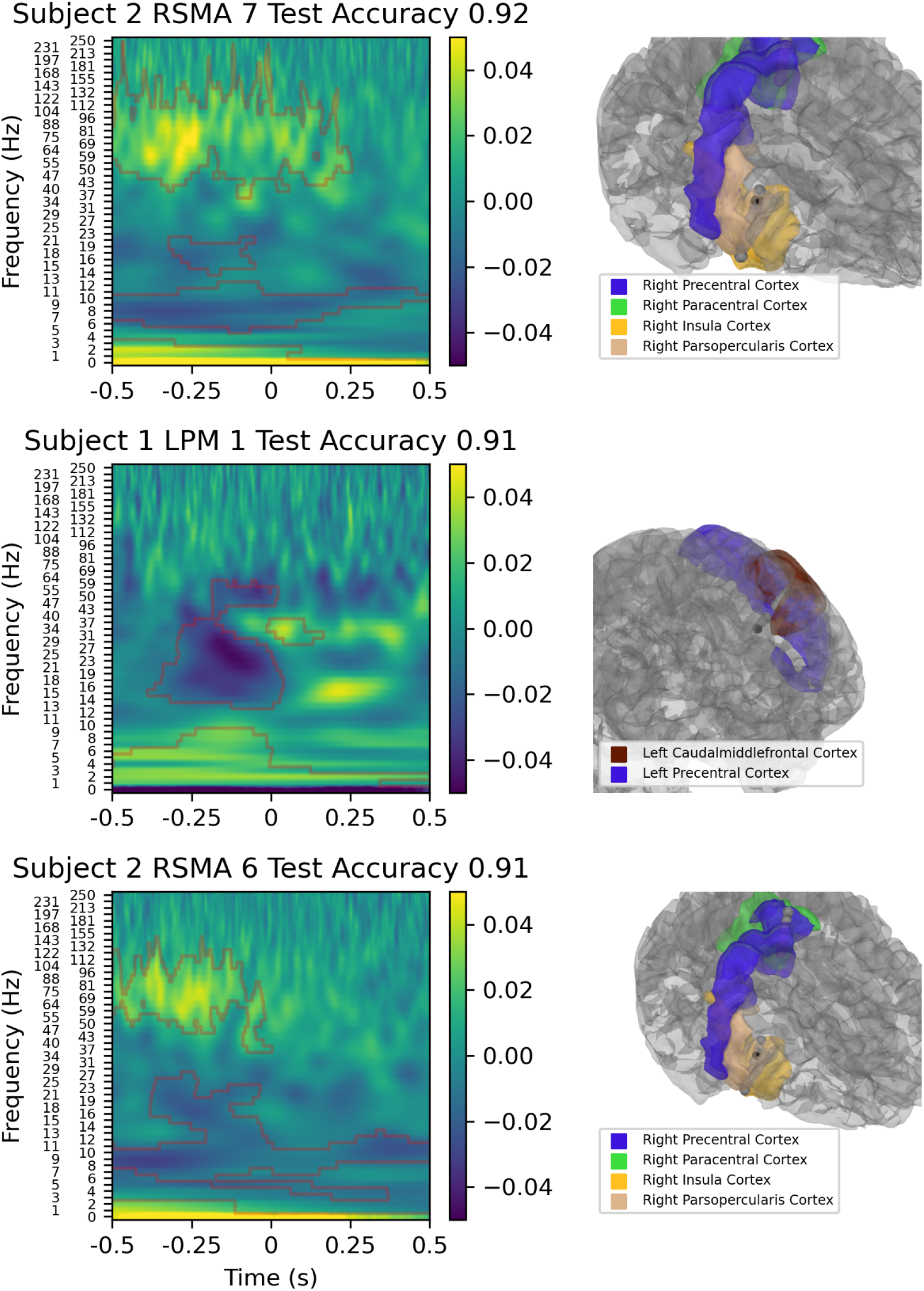
Electrode contacts with the highest classification accuracies: SVM coefficients from the spectrogram classification are shown on the left with red contours from the cluster permutation analysis, and the anatomical location is on the right with the contact colored darker than the rest of the electrode. Note that LPM 1 is located in white matter and only gray matter structures are shown.

**Figure 8.**
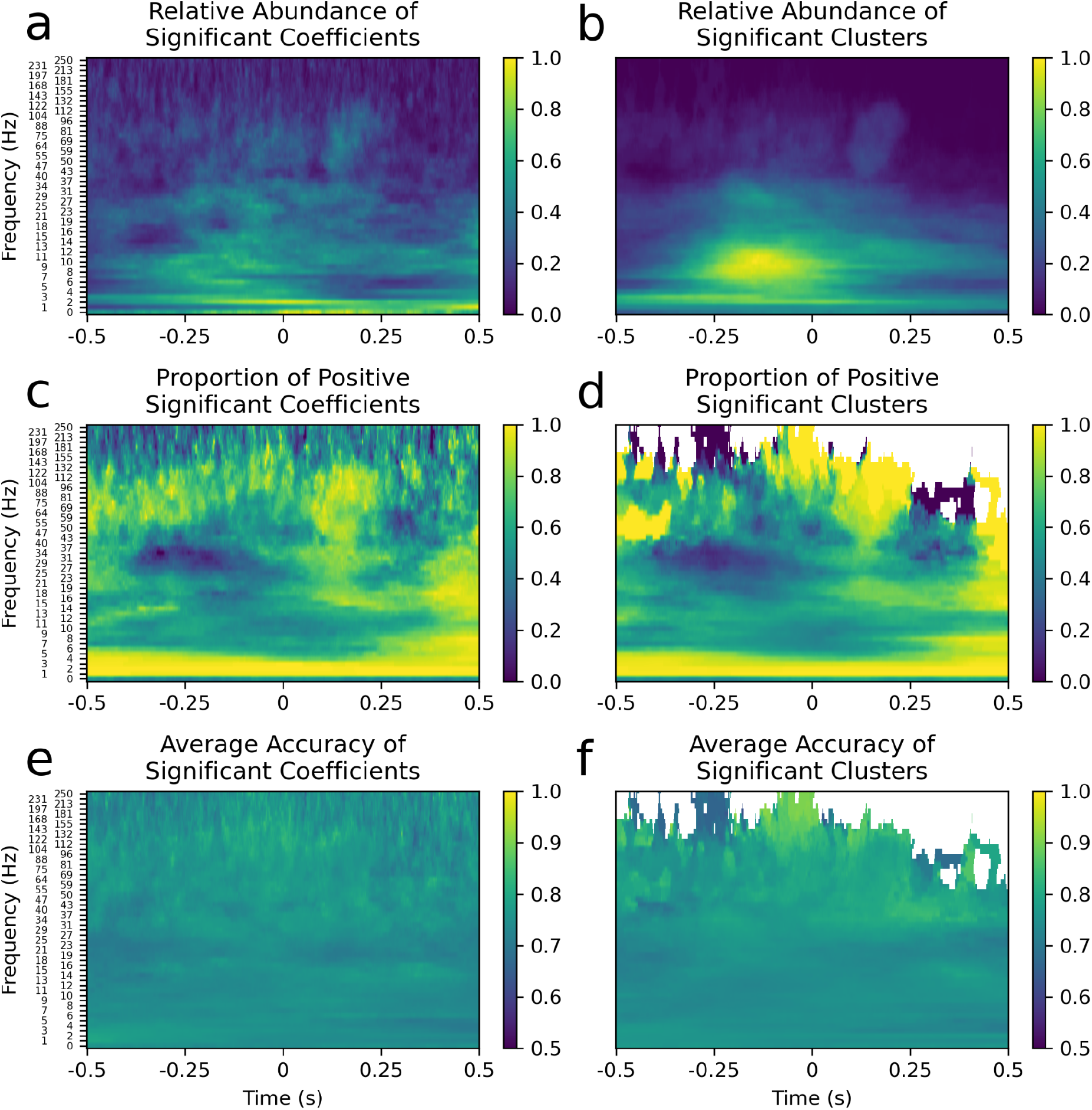
Summary feature maps of all the contacts with significant SVM classifications are shown. a and b) The relative abundance of significant SVM coefficients (a) and clusters (b); more time-frequency points tended to be significant for pre-response alpha, post-response delta or evoked potentials. c and d) The proportion of each of the significant SVM coefficients (c) and clusters (d) that were positive; pre-response beta coefficients tended to be negative, post-response beta and gamma tended to be positive and all delta tended to be positive very strongly. e and f) The average classification accuracy for significant SVM coefficients (e) and clusters (f); classifications with significant time-frequency points tended to have the same accuracies regardless of the time-frequency of the point. The cluster permutation analysis (right column) corresponded nearly perfectly to the SVM coefficient matrices (left column).

The SVM classification was used to determine which features of the spectrograms were important for classification to elucidate the mechanism of high-accuracy classifications. For each contact, the linear hyperplane optimized by the SVM was represented as a coefficient matrix that was used to evaluate the input spectrogram. The SVM classification worked by pointwise multiplying the coefficient matrix with the input spectrogram (represented in dimensionally reduced form by its PCA components), and if the sum is greater than zero the spectrogram was classified as being during movement. Thus, examining the coefficient matrix allowed us to explore which features of the spectrogram and the oscillatory patterns that they represented were related to high-accuracy classifications of movement. The coefficient matrices were assigned *p*-values for each time-frequency point using the distribution of coefficient matrices from the null classification with an *alpha*=0.01 (uncorrected) threshold.

The three best-classifying contacts using the SVM are shown in *Figure 7* to highlight areas with high classification potential and the differences between the oscillatory patterns that yielded high accuracies in different brain areas. The first contact, RSMA 7, had a positive event-related potential, a decrease in alpha power before and after the movement and an increase in gamma power before the movement. This contact is plotted in black and was in the right insular cortex. The second contact, LPM 1, had a beta desynchronization and had a large negative event-related potential that was used to classify movement. Note that the event-related potential, labeled 0 Hz at the bottom of the spectrogram, was highpass filtered at 0.1 Hz so this is not the direct current offset. Interestingly, this contact was in white matter near the left precentral gyrus. The third contact, RSMA 6, was next to the first contact, RSMA 7, and used similar features in its classification. Overall, contacts with high classification accuracies used different spectral features to classify based on their anatomical location. The anatomical locations of different spectral features are explored in the subsequent analyses.

To interpret the coefficient matrices of the SVM classifications, and the information they convey about which oscillatory patterns were important for classification, several characteristics were computed as shown in *Figure 8*. The relative abundance of significant time-frequency points was computed to determine which spectral features (i.e. time-frequency points) were the most widespread among the brain areas sampled in this study for contacts with significant classifications (*Figure 8a* and *8b*). The oscillatory patterns that these spectral features represent are likely to be the most widely-used frequencies of movement neural circuits. The proportion of positive significant coefficients (*Figure 8c* and *8d*) was computed to determine the most consistent directional patterns across brain areas. Time-frequency points that consistently had decreased power in movement spectrograms compared to baseline spectrograms are closer to zero and shown in blue. This measure was used to discriminate oscillatory patterns by their directionality; beta desynchronization, beta rebound and gamma power increase all had a consistent directionality. Interestingly, alpha power increased in some brain areas and decreased in others. Lastly, for each time-frequency point, the average accuracy was computed for all classifications in which that point was significant (*Figure 8e* and *8f*). Interestingly, all of the oscillatory patterns had remarkably similar classification accuracies. The SVM coefficient matrices (*Figure 8*, left column) were compared with a cluster permutation analysis (*Figure 8*, right column); the significant time-frequency points were in almost exact correspondence between the two analyses. This was an important validation because the determining significant time-frequency points from the SVM coefficient matrices used an uncorrected significance level with multiple comparisons.

To analyze the anatomical locations of oscillatory patterns, important spectral features were selected by inspection based on the feature maps in *Figure 8*. The features chosen were the event-related potential, a movement delta increase before the movement but extending to slightly after, a beta decrease immediately before movement, both a high- and low-beta increase after movement and a gamma increase immediately after movement. These band-based categories are highlighted by a red box surrounding the time-frequency area of a relevant feature map in the first column *Figure 9*. The second column shows the distribution of significant time-frequency points relative to the area of the red box, where -1 represents when all points within the box are significant and negative and 1 represents when all points are significant and positive. The histogram was divided into tertiles with the negative tertile in blue and the positive tertile in yellow. The contacts corresponding to classifications that used these features are plotted in the final three columns to show their position on the template brain. In addition, the oscillatory patterns of interest are shown in a different format in *Figure 10*; anatomical labels from the Desikan-Killiany atlas for the individual patient are used to highlight brain structures related to each oscillatory feature. Each feature was observed to have a specific distribution and may be a single motor network communicating via that particular oscillatory pattern. These oscillatory patterns and putative motor networks are very likely interrelated but each may have its own specific functional relevance.

**Figure 9.**
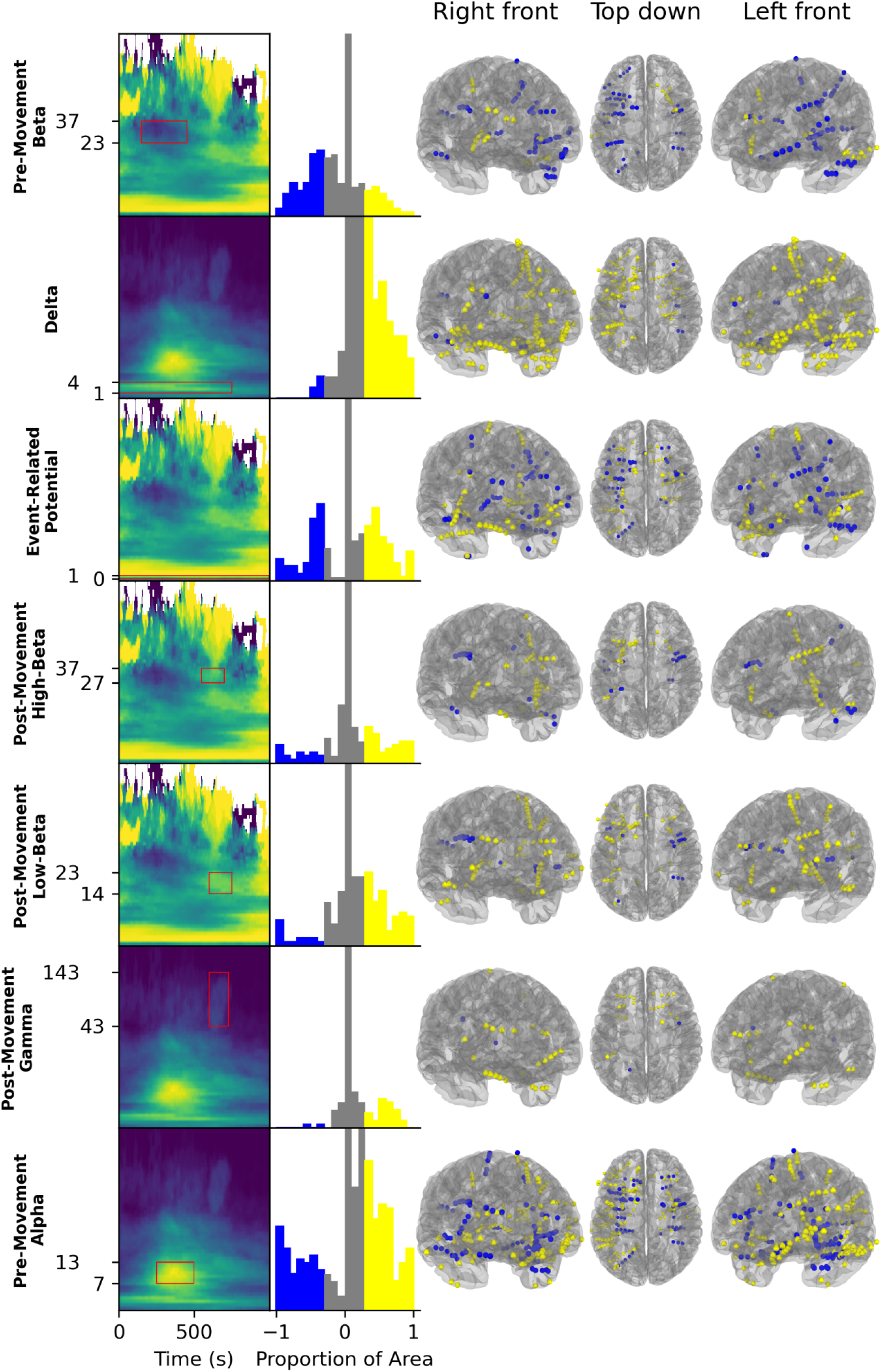
Anatomical locations of spectral features are shown on the template brain. In the first column, the time-frequency period in consideration is surrounded by a red box over a cluster permutation analysis feature map (Figure 8). The choice of which time-frequency areas to consider was made based on the relative abundance feature map and the proportion of positive clusters feature map, but only the feature map that shows the pattern more clearly is shown. The second column has histograms of the proportion of the selected area in a significant cluster for each contact, with -1 being all points in a significant negative cluster and +1 being all points in a significant positive cluster. The negative tertile is colored blue and the positive tertile is colored yellow. Note that the vast majority of contacts are near zero so the y-axis is truncated to show the rest of the distribution. The final three columns show the anatomical locations of contacts with each feature (i.e. in the negative and positive tertiles), warped to a template brain.

**Figure 10.**
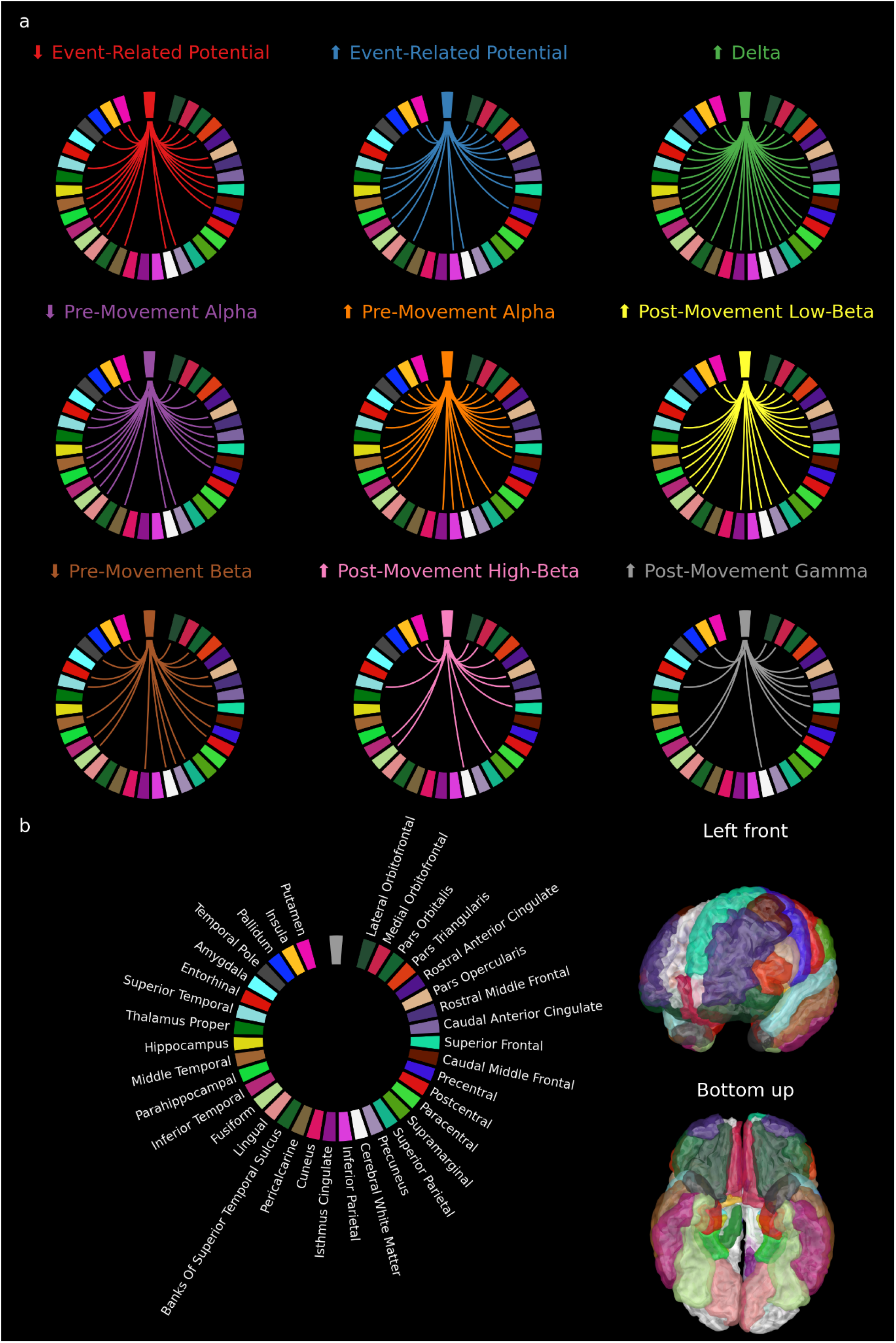
a) The Deskian-Killiany atlas labels within 3 mm of each contact with a significant-accuracy SVM classification that used that oscillatory pattern as determined in Figure 9. b) The labels used in (a) and their locations on the template atlas. Note that the atlas labels are sorted clockwise by their angle in the sagittal plane, starting with frontal areas and wrapping around to temporal areas.

## Discussion

Classification of sEEG data during periods of movement with an SVM presents a way to leverage the power of machine learning while maintaining information about what contributes most to classification. We found that many electrode contacts had high-accuracy movement-related classifications throughout the brain; not all patients had contacts implanted in sensorimotor areas but nearly 50% of all contacts had classifications unlikely to be observed by chance. The contacts with above-chance accuracy are putatively in areas of distributed motor networks, or at least part of networks that were heavily recruited during a simple motor task. The locations of these contacts replicate and extend previous results in ECoG and EEG; beta desynchronization was observed in sensorimotor areas and the inferior frontal gyrus (Swann et al., 2009) and extended to more superior frontal areas than previously described. Beta-gamma phase-amplitude coupling (PAC) has been observed to be abnormally elevated in Parkinson’s disease (de Hemptinne et al., 2015) so understanding the full spatial extent of the beta oscillations that are aberrantly coupled to gamma could inform the development of medical interventions that manipulate this neural circuit. Alpha power modulations were observed in prefrontal, sensorimotor and mid-temporal regions, as were previously reported, but extended to subcortical and parietal areas that have not been linked with alpha. The areas that have not been associated with alpha rhythms may help link the many functional correlates of the alpha/mu oscillation (Pineda, 2005). The positive and negative event-related potentials were widely distributed in frontal, parietal and superior temporal areas. Somatosensory-related potentials have been successfully modeled by increases in excitatory output from the granular layer of somatosensory cortex (Jones et al., 2007). Interestingly, gamma power, which has also been associated with a greater balance of excitation compared to inhibition, was observed to have a different distribution with greater frontal and less temporal activation. Gamma has been correlated with single-unit spiking, and tends to be biased towards recording large excitatory neurons (Manning et al., 2009). The timing of gamma compared to the event-related potentials and their observed distributions suggest different roles in motor networks, and that they are able to measure excitatory neurons differently, perhaps with event-related potentials being more related to phase resetting (Sauseng et al., 2007). Lastly, the increase in delta power in almost all recorded brain areas during the peri-movement period may be related to an increase in concentration during the task (Harmony, 2013). In general, the relative timing of increases and decreases in oscillatory power relative to movement provides insight into how oscillations might travel between brain regions and consequently how information might flow.

Previous studies that have attempted to decode patterns in sEEG data related to movement without machine learning have had mixed results. Attempts to decode movement direction (Johnson et al., 2017) and path information (Breault et al., 2017) have found few modulated areas and correlations that reached statistical significance but were on the order of *r*=0.2. A study decoding movement speed that used a more complex classification method, least absolute shrinkage and selection operator (LASSO) linear regression, had a higher correlation of *r=*0.4 on average but with a large range of correlations across subjects and a mean squared error greater than one (Breault et al., 2019). Unlike these studies, we chose to classify using a baseline without movement to quantify the extent to which the brain areas recorded by sEEG were related to movement in general and focused on individual electrode contacts instead of patient-level data. The greater contrast between the baseline and movement conditions, and having only these two conditions, increased the rate of learning so that the classifier could be more successful with fewer training trials. Additionally, since the majority of sEEG electrodes were outside primary motor areas, classifying the combined effect from all movement-related activity is appropriate for characterizing the extent of all movement-related brain networks. Focusing on a specific property of movement, such as speed, depends on sampling more specialized motor networks which may not be extensively covered by the sEEG montages of the patients in the study. By using a simpler comparison, focusing on movement more generally, and using machine learning, we described the extent of brain networks related to movement beyond what had been reported in previous studies.

The primary limitations of this study are the lack of a consensus for methodological choices for sEEG and the simplicity of the machine learning approach. Using an average reference allowed us to study the activity at individual channels (compared to a bipolar referencing scheme where activity is localized between two contacts) but was dependent on the rest of the recording montage which was unique to every patient. However, with over 100 channels per montage, the average reference is likely to be stable and reproducible given a similar sized montage. All of the patients in this study had similar power spectral density of their sEEG data, which supports this claim. One potential issue with an average reference is the introduction of an event-related potential from the average reference. When this analysis was rerun with a bipolar referencing scheme, a similar number of contacts had significant classifications relative to the null distribution, which is evidence that this is not the case. Further methodological work will clarify the optimal referencing scheme and other methodological choices. Additionally, we chose a machine learning method that prioritized interpretability, which came at the cost of classification accuracy; a more complex algorithm could have fit the data better. Future work may use more powerful machine learning approaches which will likely achieve better accuracies but would very likely have less interpretability compared to this analysis.

The results of this analysis suggest several further avenues of inquiry related to the use of sEEG data to decode movement brain networks. One future direction would be to use spectral connectivity for classification instead of spectrograms. This would test whether brain areas that have the same oscillatory patterns are part of the same neural circuit or whether there are multiple independent circuits that operate at the same oscillatory frequencies. Another direction that merits further study is the feasibility of adding anatomical targets from our results to control a brain-computer interface (BCI). Patients with motor deficits are often willing to undergo invasive surgery to restore function with a BCI. Since ECoG has not yet been able to fully restore movement (Miller et al., 2020) and since sEEG uses minor 2.4 mm bolt holes, which are less invasive than the craniotomy required for ECoG, improving BCI control by adding sEEG to sample different motor network nodes is an option that should be considered. Lastly, our results are proof-of-concept that intraoperative functional brain mapping could be more efficient with machine learning. Conventional passive (non-stimulation) mapping of functionally significant motor areas uses modulation of a predefined frequency band, commonly gamma (Kreidenhuber et al., 2019). Using an SVM or similar spectrogram-based classification would be more accurate because it includes a wider range of frequencies, including and extending beyond the predefined band.

This analysis also provides evidence for the power and reliability of a machine learning classification of electrophysiology data, specifically sEEG data. Stereo-EEG is implanted for clinical indications in brain areas that are sparsely and idiosyncratically covered, sampling a large range of brain networks that perform many different functions. Combining the information from isolated electrodes in sEEG recordings within and across patients into interpretable networks without the results being confounded by the cacophony of other networks is a complicated task. Our results provide preliminary evidence that a machine learning approach may be more fruitful at decoding this complicated information; previous studies often focused on one oscillatory pattern at a time, whereas here, with only eight patients, all of the most widely-reported oscillatory patterns were replicated.

Finally, we observed a normal distribution of sEEG contact classification accuracies, rather than a bimodal pattern that would correspond to movement-related and non-movement related contacts. This suggests that either different patterns of motor network activation are occurring on each trial, leading to missed classifications, or that only some of the population of neurons sampled by each sEEG electrode contact are involved in movement. Previous work has shown that, at a cellular level, stable sequences of activity are observed during simple motor tasks (Recanatesi et al., 2022). Thus, the more likely explanation is that a large portion of the population of neurons recorded by an sEEG channel are not involved in movement, causing classifications to fail on trials where the activity of movement-unrelated neural circuits is more prominent. Thus, this study provides evidence that movement neural circuitry is distributed such that there is a continuum from relatively homogenous sensorimotor-dominated areas, like primary motor cortex, to areas where only a minority of neurons are modulated by movement.

Overall, we identified distinct time-frequency patterns with high-accuracy classifications using electrophysiological recordings of movement. Contacts with accurate classification are more widespread in anatomical location than previously described. The specific structural networks that communicate with these oscillations have yet to be fully determined but this characterization makes substantial progress toward that goal.

## Acknowledgements

We would like to acknowledge Preeya Khanna for her helpful feedback on the manuscript. This work is supported by the Renée James Seed grant to Accelerate Scientific Research from University of Oregon (APR, NCS; PI: N. Swann) and NIH-1RF1MH117155-01 (BS, AM, AMR, PI: A. Raslan).

